# CANT-HYD: A curated database of phylogeny-derived Hidden Markov Models for annotation of marker genes involved in hydrocarbon degradation

**DOI:** 10.1101/2021.06.10.447808

**Authors:** Varada Khot, Jackie Zorz, Daniel A. Gittins, Anirban Chakraborty, Emma Bell, María A. Bautista, Alexandre J. Paquette, Alyse K. Hawley, Breda Novotnik, Casey R. J. Hubert, Marc Strous, Srijak Bhatnagar

**Affiliations:** Department of Geoscience, University of Calgary, Calgary, Alberta, T2M 4G5, Canada; Department of Biological Sciences, University of Calgary, Calgary, Alberta, T2M 4G5, Canada

## Abstract

Discovery of microbial hydrocarbon degradation pathways has traditionally relied on laboratory isolation and characterization of microorganisms. Although many metabolic pathways for hydrocarbon degradation have been discovered, the absence of tools dedicated to their annotation makes it difficult to identify the relevant genes and predict the hydrocarbon degradation potential of microbial genomes and metagenomes. Furthermore, sequence homology between hydrocarbon degradation genes and genes with other functions often results in misannotation. A tool that systematically identifies hydrocarbon metabolic potential is therefore needed. We present the Calgary approach to ANnoTating HYDrocarbon degradation genes (CANT-HYD), a database containing HMMs of 37 marker genes involved in anaerobic and aerobic degradation pathways of aliphatic and aromatic hydrocarbons. Using this database, we show that hydrocarbon metabolic potential is widespread in the tree of life and identify understudied or overlooked hydrocarbon degradation potential in many phyla. We also demonstrate scalability by analyzing large metagenomic datasets for the prediction of hydrocarbon utilization in diverse environments. To the best of our knowledge, CANT-HYD is the first comprehensive tool for robust and accurate identification of marker genes associated with aerobic and anaerobic hydrocarbon degradation.

## INTRODUCTION

Hydrocarbons are diverse compounds consisting of carbon and hydrogen atoms that differ in size, structure, and reactivity. They can be the product of geological processes as well as produced biogenically by organisms in all domains of life (1–3). Assessing hydrocarbon use by microorganisms, as a source of carbon and/or energy, is important for evaluating the consequences of hydrocarbon presence or contamination (4), understanding the global carbon cycle (5), and for industrial applications, such as the synthesis of biocatalysts (6). Degradation of hydrocarbon molecules is kinetically challenging due to the chemical inertness of the organic C-H bond, and when present, the stability of aromatic ring structures (7). Microorganisms employ a range of enzymes to use hydrocarbons (7, 8) in oxic and anoxic conditions. Catabolism of these hydrocarbons is coupled with reduction of terminal electron acceptors such as oxygen, nitrate, sulfate, and iron or via syntrophy with methanogens (9).

The discovery of hydrocarbon degrading microorganisms has traditionally relied on cultivation in the laboratory using hydrocarbon substrates (10, 11). Successful cultivation preceded the identification of genes involved in hydrocarbon metabolism with techniques such as gene knockouts, protein expression analyses, and gene sequencing (12–17). These studies are crucial for providing fundamental knowledge on the ever-growing diversity of hydrocarbon degrading microorganisms as well as uncovering new degradation pathways. The recent exponential rise in sequence data and the consequential increase in known microbial diversity have provided new opportunities to explore hydrocarbon degrading potential in diverse environments and uncultured microorganisms. One approach for exploring sequence data is to annotate genes using Hidden Markov Models (HMM). HMMs are trained on the multiple sequence alignments of amino acid sequences and produce position-specific scores and penalties when searching query sequences. HMMs have better sensitivity and recall for identifying homologs of conserved protein domains, compared to conventional pairwise alignment tools such as blastp (18), which use a position-independent scoring matrix (19). Detection of metabolic potential in whole genomes or metagenomic datasets is generally accomplished using functional annotation tools aided by HMM databases such as KEGG (20) and Pfam (21). While these large databases can confidently identify central metabolic and other well studied pathways, specific HMMs and tools for accurate annotation of catalytic genes in hydrocarbon degradation pathways are currently lacking. Genes involved in hydrocarbon degradation can share sequence similarity to genes from other metabolic pathways and consequently, are often misannotated (22, 23). Hence, there is a need for a purpose-built tool for the accurate detection of hydrocarbon degradation pathways in sequence data.

Here we present the Calgary approach to ANnoTating HYDrocarbon degradation genes (CANT-HYD), a database containing 37 HMMs designed for the identification and annotation of marker genes that are critical for the aerobic and anaerobic degradation of alkane and aromatic hydrocarbons. CANT-HYD is tested and validated against more than 70 genomes of known hydrocarbon degrading bacteria, representing a broad spectrum of hydrocarbon metabolism. Using these validated HMMs, over 30,000 representative genomes covering the entire bacterial and archaeal tree of life are analyzed to identify hydrocarbon degrading microorganisms. Forty-one publicly available metagenomes from diverse environments are also analyzed using CANT-HYD to explore hydrocarbon degradation potential in diverse environments.

## METHODS

### Selection and clustering of archetype reference sequences

Enzymes involved in the activation of hydrocarbon substrates in aerobic and anaerobic hydrocarbon degradation pathways of aliphatic and aromatic compounds were identified through a literature search (Figure 2). Amino acid sequences encoding the catalytic subunits of these enzymes were obtained from Genbank and were classified as either ‘experimentally verified’ or ‘putative’. ‘Experimentally verified’ sequences refer to those from published studies with experimental proof of the intended function. Experimental proof consisted of gene cloning or protein purification and corresponding enzyme assays, or gene knockout studies. Gene sequences labeled ‘putative’ refer to sequences with strong evidence of function but lacking these definitive prerequisite analyses. Putative sequences often originated from isolates or enrichment cultures where there is evidence of hydrocarbon degradation or genomic and/or proteomic evidence for the enzyme responsible. The resulting curated 105 ‘archetype’ gene sequences from 53 different species (Supplementary Table ST2) were clustered into homologous groups based on ≥ 20% amino acid identity as determined by blastp v2.9.0 (18). Sequences that did not cluster at 20% were either manually added into groups of similar function or left as singletons.

**Figure 1.**
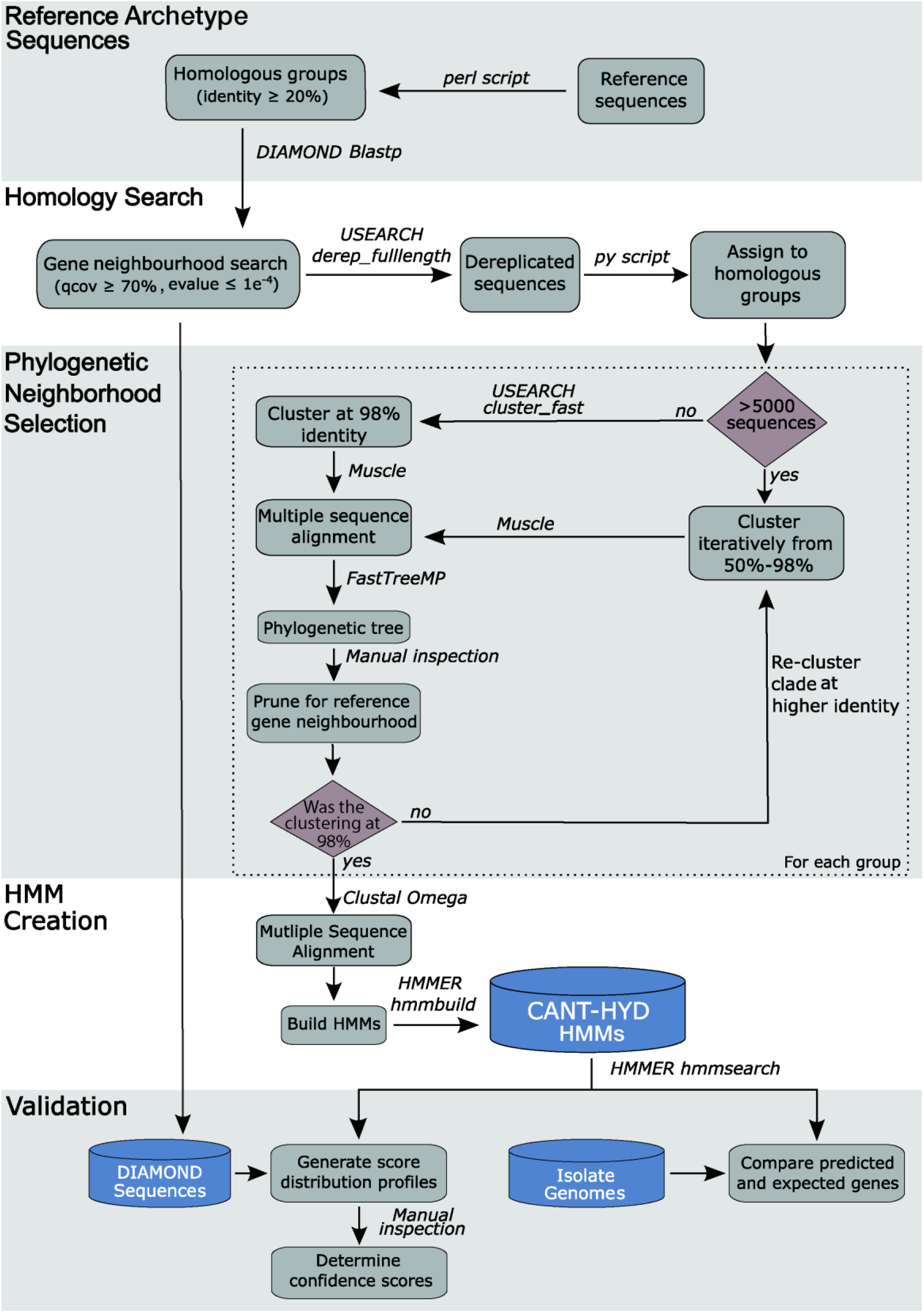
Workflow of the process underlying the creation of the CANT-HYD HMM Database.

**Figure 2.**
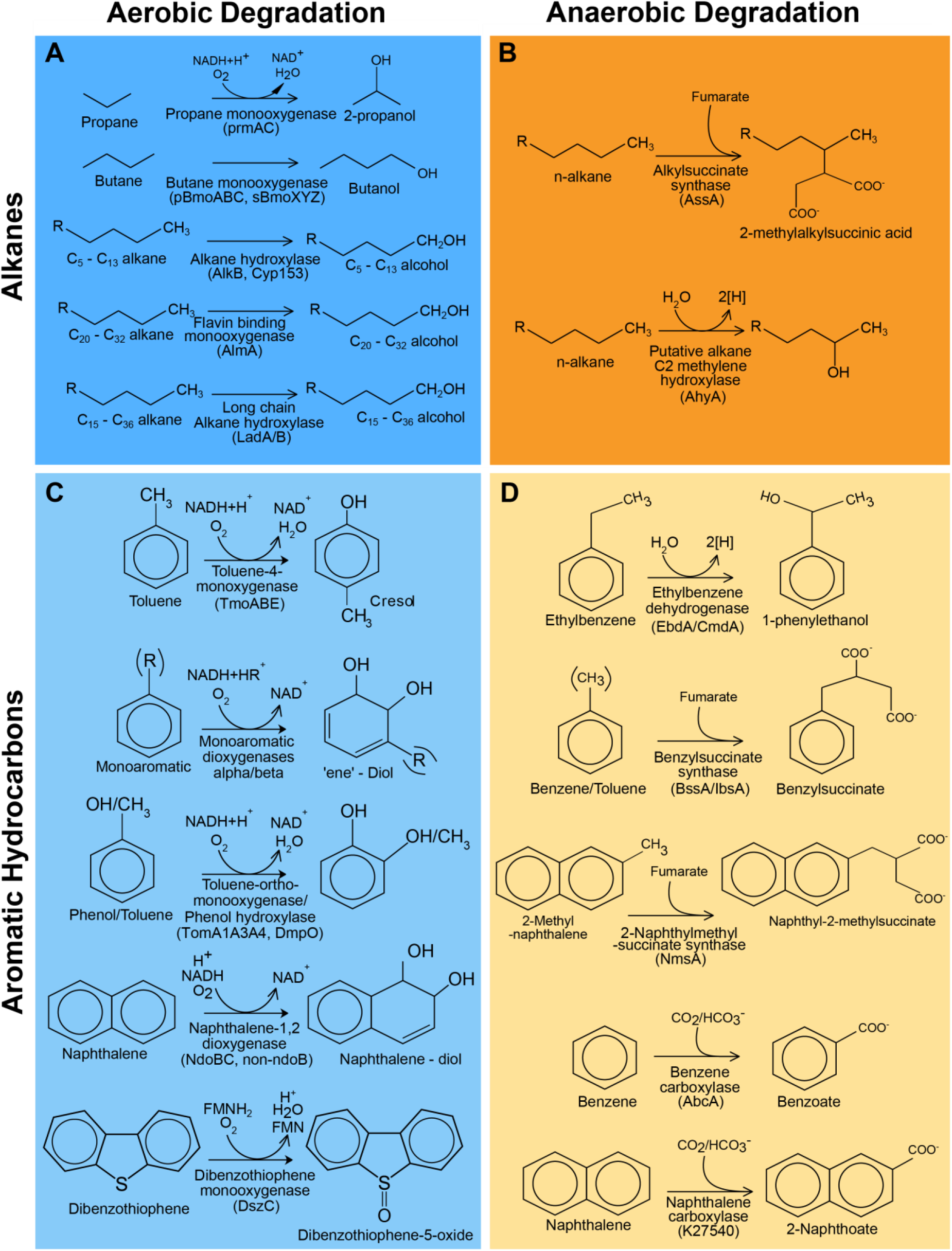
Hydrocarbon degradation reactions covered by CANT-HYD. Reactions for the degradation of alkanes through aerobic (**A**) and anaerobic pathways (**B**) and degradation of aromatic hydrocarbons through aerobic (**C**) and anaerobic (**D**) pathways.

### Homology search to obtain genes similar to experimentally verified and putative archetypes

Genes sharing sequence homology to the 105 genes with experimental support were recruited from the NCBI non-redundant (nr) protein database using a Diamond homology search (24). All hits with query coverage ≥70% and e-value ≤ 10^-4^ were retained and assigned to the query’s homologous group. The sequences were dereplicated, followed by clustering at 98% amino acid identity using the USEARCH v9.0.2 *derep_fulllength* and *cluster_fast* commands (25).

### Grouping genes with similar functions using phylogenetic analysis

A multiple sequence alignment was generated for each dereplicated homologous group using MUSCLE v3.8.31 (26). The alignments were used to create maximum-likelihood trees using FastTreeMP v2.1 (27) with the parameters –*pseudo* and -*spr 4*. Trees were manually inspected using iTOL v5.6.3 (28) or Dendroscope v3.6.3 (29) to identify monophyletic clades of genes containing experimentally verified archetype sequences. (See Supplementary Figure S1). Sequences within a given monophyletic clade were extracted and used as seed sequences for the respective HMM.

### Processing homologous groups with >5,000 sequences

For any homologous group where clustering at 98% amino acid identity resulted in >5,000 representative sequences, a nested clustering, and phylogenetic pruning approach was implemented to overcome computational challenges. The homologous group was first clustered at a lower identity (e.g., 50%), followed by alignment, phylogenetic reconstruction, and phylogenetic neighborhood pruning as described above. The process of pruning the phylogenetic neighborhood of the reference sequence(s) was iterated until the prune group was ≤ 5,000 sequences or the clustering identity was raised to 98% (Figure 1), at which point the seed sequences for HMM were selected as described above.

### HMM creation and determination of cutoff scores

Each gene set was aligned using Clustal-Omega v1.2.4 (30) followed by manual inspection of the alignment using Jalview v2.11.1.4 (31), and generation of HMMs using the *hmmbuild* command of HMMER v3.2.1 (32). The HMMs were used to search the archetype reference sequences and ‘DIAMOND Sequences’ database (Figure 1) using *hmmsearch* of HMMER v3.2.1 (32). The domain scores for each HMM were plotted to visualize the frequency distribution pattern of the scores (Supplementary Figure S2). A ‘trusted’ and a ‘noise’ cutoff was chosen for each HMM using the score distributions. The trusted cutoff is the domain score above which a sequence can be confidently annotated for the function, as all experimentally verified genes used for the HMM scored above this cutoff. The noise cutoff was chosen to exclude genes with some sequence homology but potentially a different function. Hits scoring below the noise cutoffs are expected to not carry out the function represented by the HMM. Supplementary table ST1 includes information on genes that are the closest phylogenetic relatives to the archetype sequences of CANT-HYD HMMs but do not play a role in hydrocarbon degradation.

### Validation of CANT-HYD HMMs using genomes of known hydrocarbon degraders

Seventy-two genomes of microorganisms with published experimental evidence of an ability to degrade hydrocarbons were downloaded from Genbank and Refseq (33) and categorized by the type of substrate and respiration. If the exact strain was not available, its closest relative from GTDB was chosen. For example, *Aromatoleum aromaticum* EbN1 anaerobically degrades aromatic compounds (34). The genomes were searched using CANT-HYD HMMs and the resulting gene annotations, scoring above the trusted cutoff, were compared to the established degradation capability of the organism. If a gene hit multiple HMMs above the confidence threshold, it was assigned to the highest scoring HMM.

### Analysis of GTDB genomes to identify potentially novel hydrocarbon degrading bacteria

The GTDB database (05-RS95 17th July 2020) (35) of representative bacterial and archaeal genomes was downloaded and searched using the CANT-HYD HMMs. For further investigation, gene sequences from Cyanobacterial genomes with hits to LadA beta (above the noise cutoff) were combined with archetype reference sequences of long-chain alkane monooxygenases (LadA-alpha, LadA-beta, and LadB). The combined sequences were then clustered at 70% amino acid identity using USEARCH v9.0.2132_i86linux64 (25) *cluster_fast*. Representatives sequences of each cluster were then aligned using Muscle v3.8.31 (26), followed by a maximum-likelihood phylogenetic reconstruction using FastTreeMP v2.1 (27) (Supplementary Data SD1).

### Analysis of diverse metagenomes using CANT-HYD

Metagenomes representing diverse environments such as petroleum reservoirs (36–39), oil spill experimental microcosms (40, 41), marine systems (42–45), host-associated microbiomes (46–48) and other environments (49, 50), were downloaded either as unassembled raw data from the NCBI SRA or as predicted gene sequences from the JGI Genome Portal (Supplementary Table ST4). Raw reads from unassembled metagenomes were filtered using BBDuk (sourceforge.net/projects/bbmap/) for a minimum quality of 15 and a minimum read length of 150bp. Reads passing quality control were assembled using MEGAHIT (51) with default parameters, followed by gene calling by Prodigal v2.6.3 (52) with the metagenomic option (*-p meta*). The amino acid sequences of predicted genes were searched against the CANT-HYD database using the *hmmsearch* command of HMMER v3.2.1 (32) and only hits scoring above the noise cutoff for each HMM were visualized. Hit count for each metagenome was normalized by the total number of predicted genes.

## RESULTS AND DISCUSSION

### Validation of CANT-HYD HMMs

Genomes of 72 microorganisms with experimental evidence of hydrocarbon degradation were analyzed with the CANT-HYD HMMs for validation. For 62 out of 72 organisms, gene predictions using CANT-HYD were consistent with experimental data (Figure 3). Of the remaining ten genomes, two genomes had hits with a score between the noise and trusted cutoffs, and eight genomes lacked hits above the noise cutoff. In a few instances, genomes were not available for the exact strain and a GTDB representative genome was used in their place. Although GTDB representatives share 95% average nucleotide identity with the cluster they represent, hydrocarbon degradation genes may be missing or different. Three genomes isolated on phenanthrene, chlorophenol and benzoate did not yield any hits to any CANT-HYD HMMs. Although these substrates can be degraded by dioxygenases which share homology with mono- and polyaromatic ring hydroxylating dioxygenases, the lack of hits, even below the noise cutoff, indicates that the three organisms potentially use alternative metabolic pathways which were not covered by CANT-HYD. CANT-HYD predicted additional or unreported hydrocarbon substrate degradation capabilities for 16 genomes. For example, genes for toluene-2-monooxygenase (Tom) and toluene-4-monooxygenase (Tmo), and monoaromatic dioxygenase (MAH_alpha and MAH_beta) were found in the genome of *Pseudoxanthomonas spadix* BD-a59, a well-known benzene, toluene, ethylbenzene, and xylene (BTEX) degrader (53). Anaerobic hydrocarbon degradation genes were only detected in the genomes of anaerobes, further showing the prediction accuracy of CANT-HYD HMMs. Every HMM had at least one hit, except for the bacterial benzene carboxylase (AbcA_1) and toluene-benzene monooxygenase beta subunit (TmoB_BmoB). Anaerobic benzene degradation via benzene carboxylase (AbcA_1) has been identified in a single uncultured organism belonging to *Clostridia*, for which a genome is currently unavailable (54). Toluene-benzene monooxygenase beta subunit (TmoB_BmoB) is found adjacent to TmoA_BmoA gene on the *Pseudoxanthomonas spadix* BD-a59 genome with a score above the noise cutoff, which indicates that it likely is a TmoB_BmoB gene divergent to the seed sequences that were used to make the HMM. Overall, these results show that CANT-HYD reliably identifies hydrocarbon degradation marker genes and can thus be used to predict the hydrocarbon degradation potential of genomes and in metagenomes.

**Figure 3.**
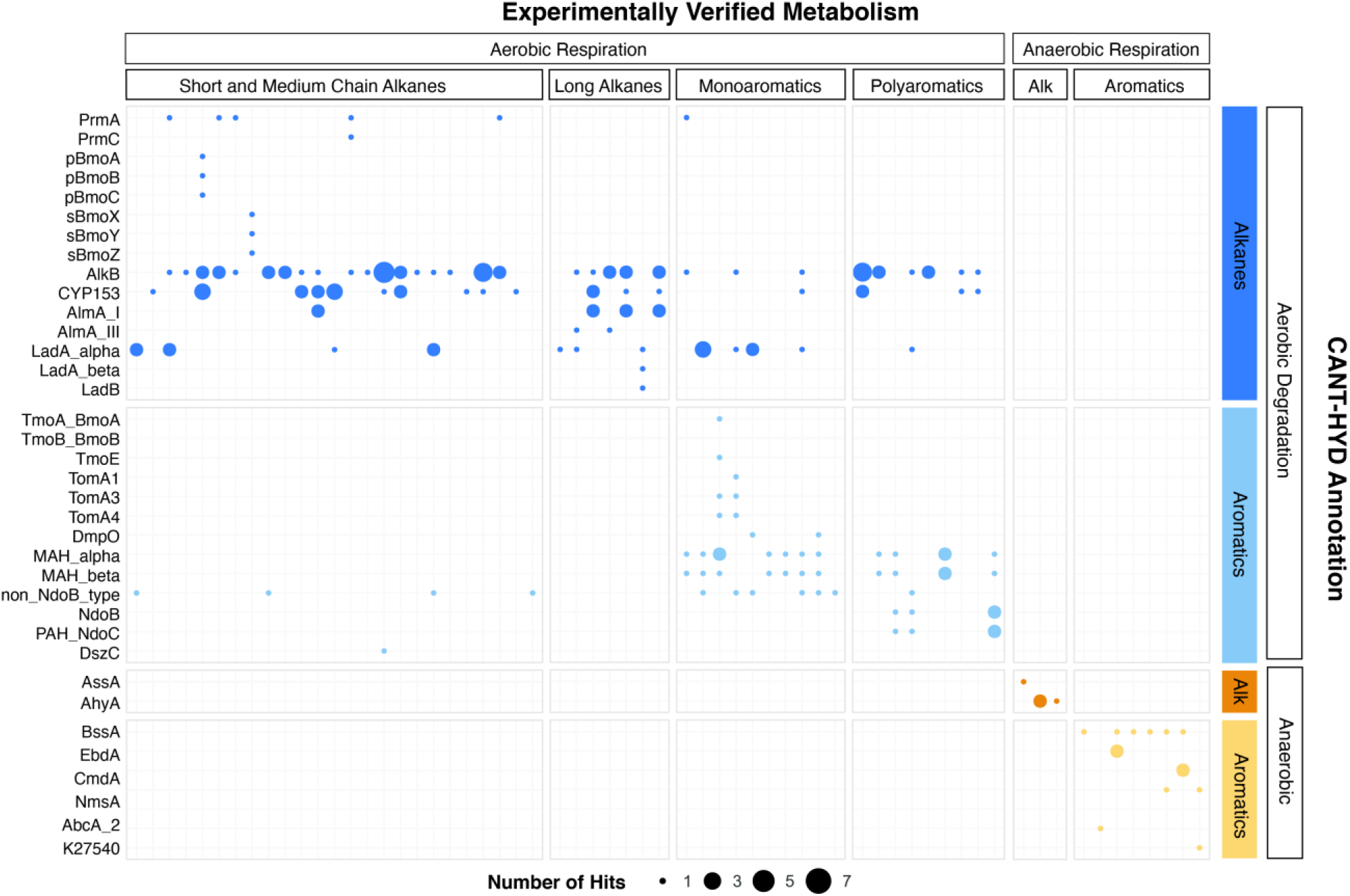
Concordance of CANT-HYD annotations of genomes and their experimentally verified hydrocarbon degradation activity. Each bubble represents hits above the confidence threshold from experientially verified hydrocarbon degrading genomes (x-axis) plotted against the HMMs of the CANT-HYD database (y-axis). The size of the bubble represents number of unique hits in the genome. The genomes (x-axis) and the CANT-HYD HMMs (y-axis) are organized by the hydrocarbon substrate and respiration. A complete list of isolate genomes, their Genbank accession, and their published hydrocarbon degradation capability can be found in Supplementary Table ST3.

### Diversity of hydrocarbon degrading bacteria and archaea

A large number of bacterial (30,238) and archaeal (1,672) genomes from the Genome Taxonomy Database (GTDB) were searched against the CANT-HYD HMMs (35). In total, 4,601 representative genomes from 18 bacterial phyla, had at least one hit to an HMM that scored higher than the trusted cutoff (Supplementary Table ST1), and in total, 5,845 genomes from 24 bacterial phyla had hits to at least one CANT-HYD HMM above the noise cutoff (Figure 4). HMM hits from diverse bacterial phyla demonstrate the widespread potential for hydrocarbon degradation across bacteria (Figure 4A and B).

**Figure 4.**
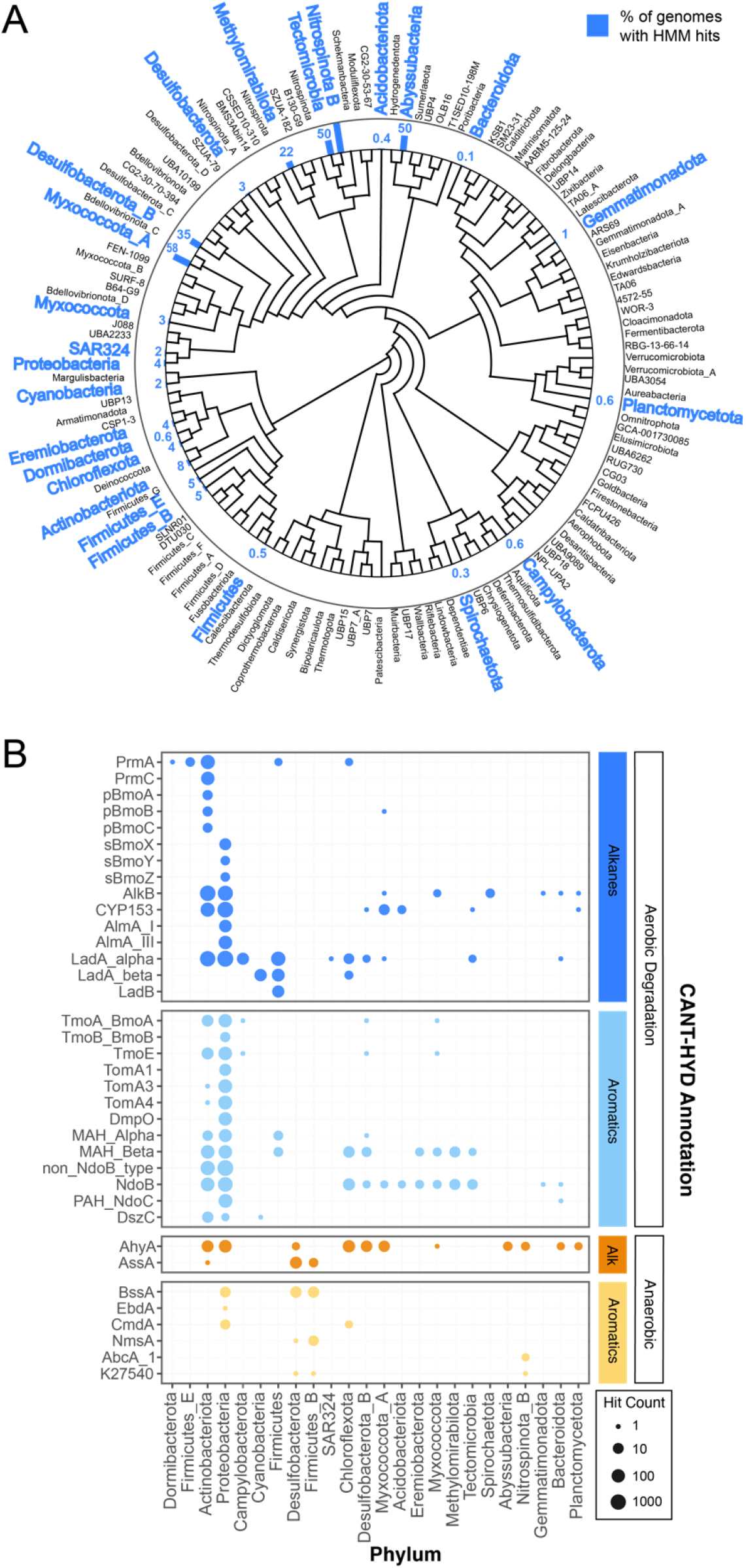
Diversity of hydrocarbon-degrading bacteria and archaea. GTDB representative genomes with CANT-HYD HMM hits shown on **A**) the GTDB phylogenomic tree collapsed at phylum level. Phyla with genomes containing genes with a CANT-HYD HMM score greater than the noise cutoff are shown in blue. The corresponding bar and number indicates the percentage of the representative genomes in that phylum containing at least one hit to a CANT-HYD HMM. For example, Abyssubacteria contained two genome representatives in GTDB, and one of the genomes contained a high confidence match to a CANT-HYD HMM; thus, the bar shows 50%. **B**) Distribution and number of hits for each phylum in GTDB across the CANT-HYD HMMs.

Many of these phyla contain no cultured representatives, and therefore annotation tools like CANT-HYD become important for offering clues about their metabolic potential. For instance, the potential for hydrocarbon degradation was found in genomes of poorly represented phyla including Abyssubacteria, Tectomicrobia, and Eremiobacterota (Figure 4B, Supplementary Figure S4). The phylum Abyssubacteria, often found in association with subsurface and hydrocarbon contaminated environments (55), had hits to anaerobic alkane degradation (AhyA). Eremiobacterota (formerly WPS-2), previously found in hydrocarbon enrichment cultures (56), had three genomes with aerobic aromatic hydrocarbon degradation potential (NdoB, MAH_alpha, MAH_beta). Two genomes from Tectomicrobia had high HMM scores to enzymes responsible for the aerobic degradation of monoaromatics (MAH_Beta), polyaromatics (NdoB), and long-chain alkanes (LadA_alpha and CYP153). There is currently no literature associating Tectomicrobia with hydrocarbon containing environments, however, high confidence matches to CANT-HYD HMMs suggest that they may have a previously unidentified role in the aerobic metabolism of a range of hydrocarbons.

#### Hydrocarbon degradation in Archaea

Archaea contained fewer hydrocarbon degradation genes compared to Bacteria. Only four genomes, all from the phylum Halobacteriota, had HMM hits above the trusted cutoff. Another 102 genomes, also from Halobacteriota, had at least one HMM hit above the noise cutoff (Supplementary Figure S3). The phylum Halobacteriota (formerly a member of phylum Euryarchaeota) is known to contain halophilic hydrocarbon degrading species (57, 58). The identification of only a few archaeal hydrocarbon degraders may also be due to either a lower representation of sequenced archaeal genomes, or an increased phylogenetic distance of archaeal hydrocarbon degradation genes to the largely bacterial sequences that have been experimentally validated, or methanotrophy, the most well studied archaeal hydrocarbon degradation, is not covered by CANT-HYD. As more experimental evidence of archaeal genes emerges, the annotation of archaeal hydrocarbon degradation will improve.

#### Cyanobacteria as alkane degraders

Thirty-six genes from 29 cyanobacterial GTDB representative genomes, mostly from the family Nostocaceae and the genera *Nostoc* and *Aulosira*, were predicted to contain LadA beta, a long-chain alkane monooxygenase (Figure 5). LadA beta is one of the three ladA-type long-chain alkane monooxygenase enzymes with experimental evidence of long-chain alkane degradation (59). Phylogenetically, these cyanobacterial genes were related to the experimentally verified LadA beta sequence from *Geobacillus thermoleovorans* (BAM76372.1), suggesting that the genes perform a similar role in their photosynthetic hosts (Figure 5A). Many cyanobacterial species produce long-chain alkanes, potentially at globally relevant levels (1, 60), and alkane degradation has been observed in microbial communities with abundant Cyanobacteria (61). Thus far however, it has been inconclusive whether the alkane degradation is performed by Cyanobacteria or other heterotrophic community members, and if the Cyanobacteria are responsible, which degradation pathways they utilize (62, 63). Strong hits to LadA beta suggest that some Cyanobacterial species have the metabolic potential for long-chain alkane degradation via LadA. Experiments are needed to confirm if this genetic potential is realized by these Cyanobacteria.

**Figure 5.**
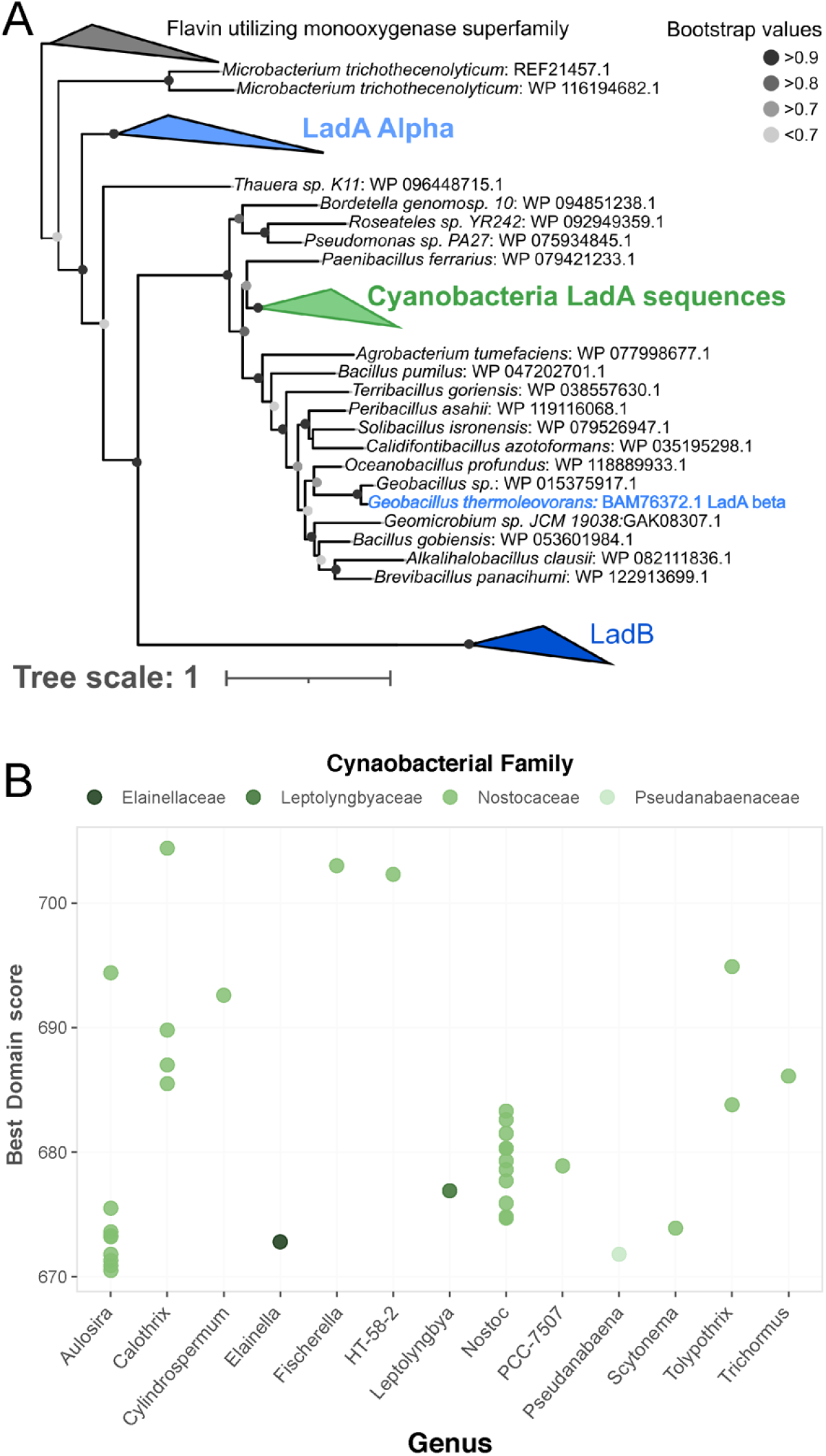
Detection of long-chain alkane monooxygenase (Lad) family of genes in Cyanobacteria. **A**) Phylogenetic tree of Lad sequences including a cluster of 102 sequences detected in Cyanobacteria genomes by CANT-HYD (green). B) Distribution of scores for LadA beta HMM hits in Cyanobacteria genomes. Original tree file for is available as Supplementary Data S1.

### Hydrocarbon degradation in diverse environments

The CANT-HYD HMMs were used to search for hydrocarbon degradation potential in 41 metagenomes, representing diverse environments including hydrocarbon degrading enrichment cultures, petroleum reservoirs, oceans, host-associated microbiomes, alkaline lakes, and hot springs. Hydrocarbon degradation genes were detected in all these environments, except for the host-associated microbiomes, which are presumed to have a limited presence of hydrocarbons (Figure 6).

**Figure 6.**
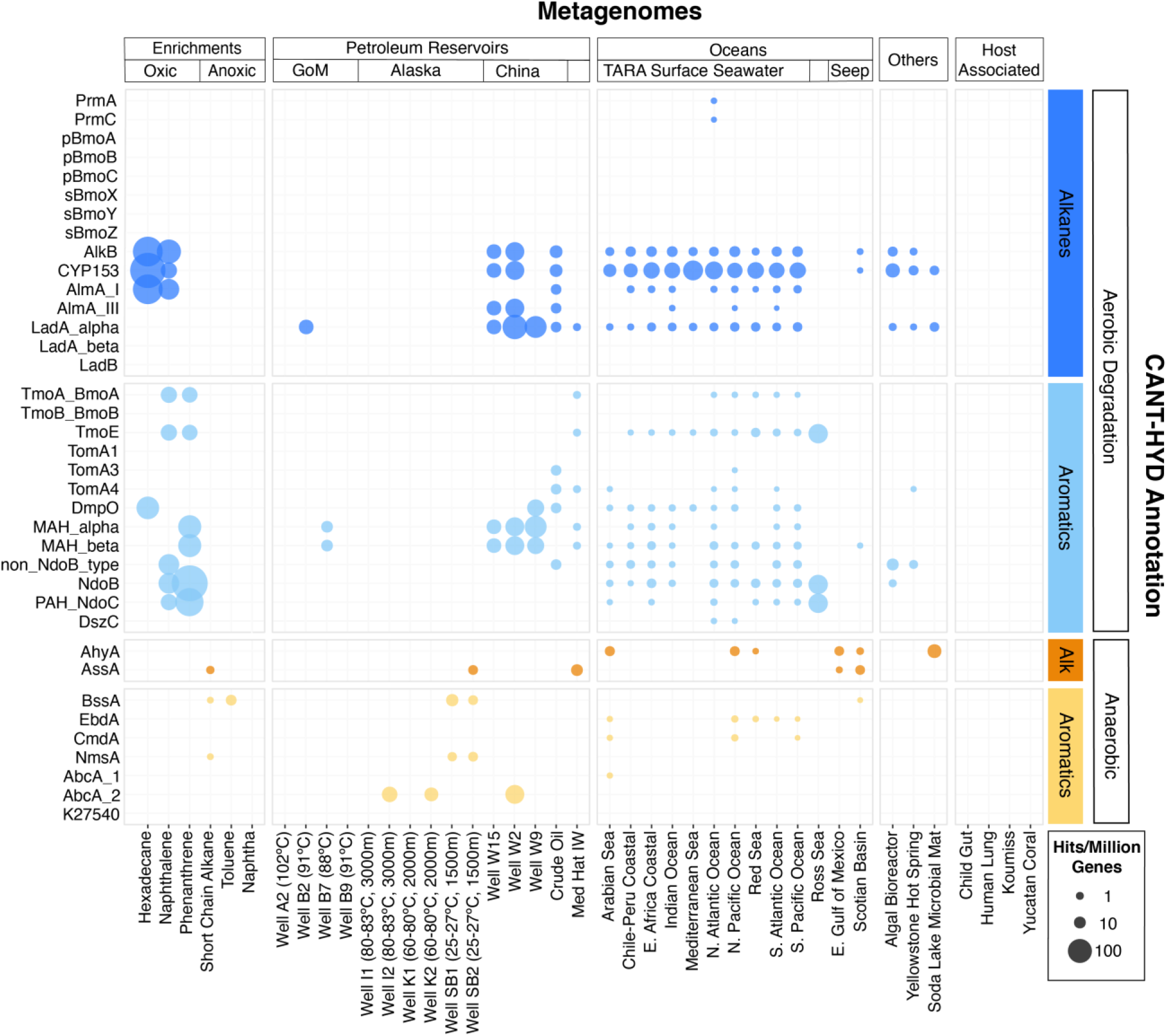
Metagenomes from diverse environments analyzed with the CANT-HYD HMMs. CANT-HYD HMM hits (scores ≥ noise cutoff) (y-axis) to each metagenome (x-axis) normalized to per million protein coding genes. The size of the bubble depicts proportion of hydrocarbon degrading genes to total genic content of the metagenome. The metagenomes (x-axis) are grouped under Enrichments (hydrocarbon-degrading enrichment cultures), Petroleum Reservoirs (produced well water from petroleum reservoirs from Alaska, Gulf of Mexico, Jiangsu and Qinghai, China, and Medicine Hat, Alberta), Oceans (cold seeps, TARA surface seawater), Host-Assoc (host-associated microbiomes), Other: (other environments). The CAN-HYD HMMs are grouped by hydrocarbon substrate (alkane and aromatic) and respiration (aerobic and anaerobic). See Supplementary Table ST4 for details regarding these metagenomes.

When normalized for total gene content, the highest proportion of hydrocarbon degradation genes were detected in the metagenomes of hydrocarbon-degrading enrichments. Genes for aerobic alkane degradation, such as AlkB, CYP153, and AlmA, were the most widely detected in this dataset. Many marker genes for aerobic hydrocarbon degradation were found ubiquitously in metagenomes from ocean surface waters, while genes for anaerobic hydrocarbon degradation were largely detected in metagenomes sequenced from anoxic habitats such as petroleum reservoirs, subseafloor sediments, and anoxic hydrocarbon degrading microcosms. Further, some degradation enzymes covered by the CANT-HYD HMMs yielded no hits, namely butane monooxygenases (pBmoA, pBmoB, and pBmoC, and sBmoX, sBmoY, sBmoZ), and benzene and naphthalene carboxylases (AbcA_1 and K27540). These HMMs were made with less than five seed sequences as they had only a few close relatives (≥ 50% sequence identity) in the public nucleotide database at the time of this work.

#### Enrichment cultures of hydrocarbon degrading microorganisms

Genes from the glycyl radical enzyme family (Ass/Bss/Nms) for anaerobic hydrocarbon degradation were identified in two out of the three anoxic cultures, namely the toluene and short-chain-alkane enrichments (40) (Figure 6). The number of genes identified in this study were half of those reported in the original, which were also annotated using custom HMMs. (Table 1). Observed discrepancies include the detection of partial gene sequences in the metagenomic assemblies (Table 1: denoted with an *) (40). Partial gene sequence matches to the CANT-HYD Ass/Bss/Nms HMMs scored below the noise cutoffs and did not pass the threshold. Partial hits to HMMs scoring below the noise cutoffs cannot be reliably annotated as they may be a partial sequence of a related gene that encodes an enzyme with a different function. In this case, the partial hits for genes encoding for glycyl radical enzymes (Ass/Bss/Nms) can be functionally similar (Supplementary Figure S1) to and share high sequence homology with pyruvate formate lyase. While CANT-HYD will not perform optimally with unassembled, partially or poorly assembled data, the noise and trusted cutoffs are recommended for a low false-positive rate, by filtering out partial genes that risk being misannotated.

**Table T1:** Number of AssA, BssA, and NmsA genes detected in this analysis compared to the original study (Tan et al., 2015). Asterisk (*) indicates partial gene homologs.

| Substrate | Short-chain alkanes |  | Toluene |  | Naphtha |  |
| --- | --- | --- | --- | --- | --- | --- |
| Genes | Original study | CANT-HYD | Original study | CANT-HYD | Original study | CANT-HYD |
| AssA | 4 + 1* | 4 | 1* | - | 1* | - |
| BssA | 3 | 1 | 1 | 1 | 1 + 2* | - |
| NmsA | 1 | 1 | - | - | 1* | - |

Several genes for aerobic hydrocarbon degradation were identified in oxic cultures inoculated with the Deepwater Horizon oil plume and enriched on hexadecane, naphthalene, and phenanthrene as hydrocarbon substrates (Figure 6). Genes for the aerobic degradation of medium and long-chain alkanes (CYP153, AlkB, and AlmA_GroupI) were detected in the hexadecane and naphthalene enrichments, while a variety of genes for the aromatic hydrocarbon degradation were detected in the oxic naphthalene and phenanthrene enrichment cultures.

#### Petroleum reservoirs and hydrocarbon biodegradation

Multiple marker genes were identified in all metagenomes from petroleum reservoirs, except for four reservoirs that were either at a high temperature (> 80°C) or were deep subsurface (Figure 6). Produced water metagenomes from petroleum reservoirs in Alaska (wells SB1, SB2, K1, K2, I1, and I2) were found to contain only anaerobic hydrocarbon degradation genes in agreement with the associated study (37). CANT-HYD further detected the presence of a putative benzene carboxylase gene (AbcA_2) in the metagenomes from two of the oil wells (K2 and I2), which was not reported in the original study. These oil reservoirs were reported to contain complete and partial genomes of a sulfide-producing archaeon, *Archaeoglobus*, and our results indicate that it can potentially metabolize benzene anaerobically.

Similar observation of an *Archaeoglobus* metagenome-assembled-genome (MAG) and its association with a benzene carboxylase (AbcA_2) gene comes from the metagenome of well W2 from the Jiangsu Oil Reservoir, China (39). Although the original study found alkyl succinate synthase genes (Ass) in an *Archaeoglobus* MAG, in this study, genes for the glycyl radical family of enzymes from these metagenomes and the *Archaeoblogus* genome scored below the noise cutoff. Any hits below the noise cutoff cannot be reliably annotated automatically and therefore require manual curation, such as using gene phylogeny to differentiate between glycyl radical enzyme and pyruvate formate lyase. A study by Khelifi et al. also shows evidence for anaerobic long-chain alkane degradation by *Archaeoglobus*, however, the genes identified as responsible for this metabolism share low sequence homology with bacterial alkyl succinate synthase alpha subunit (23) and will not be annotated by the CANT-HYD AssA HMM. Therefore, a separate HMM for archaeal alkyl succinate synthase alpha subunit would be required when strong experimental evidence for these genes becomes available. Several genes for aerobic alkane and monoaromatic hydrocarbon degradation (Figure 6) were also identified in the Well W2 metagenome, which were not originally reported. These findings highlight the utility of CANT-HYD, which can search for a comprehensive suite of hydrocarbon degradation markers, independent of *a priori* knowledge of the system.

#### Widespread hydrocarbon degradation potential in global surface seawaters

Widespread potential for aerobic hydrocarbon degradation was detected in the surface seawater metagenomes collected by the TARA Oceans survey (42, 64) and other studies [24]. Predicted hydrocarbon metabolism was largely driven by medium- and long-chain alkane hydroxylases, and ring-hydroxylating dioxygenases. This pervasive metabolic capability in the global ocean surface could be the result of biogenic alkanes synthesized by Cyanobacteria (1) or the accumulation of polyaromatic hydrocarbons by other ocean phytoplankton, resulting in a “cryptic hydrocarbon cycle” (60, 65). An ocean metagenome from the Ross Sea, Antarctica had an exceptionally high abundance of marker genes involved in polyaromatic hydrocarbon degradation. The Ross Sea is well-known for its seasonal algal blooms and rapid carbon turnover (66, 67), which have been associated with biogenic alkanes, polyaromatic hydrocarbons, and PAH degraders (1, 60, 68). Overall, our findings support the recent experimental evidence of a marine hydrocarbon cycle (60).

## CONCLUSION

Here, we describe CANT-HYD, an HMM database of marker genes for hydrocarbon degradation. This phylogenetically informed database accurately identifies 37 genes relevant to aerobic and anaerobic metabolisms of aliphatic and aromatic hydrocarbons in genomes and metagenomes. Each CANT-HYD HMM includes a manually curated trusted and noise cutoff score for automated reliable detection of these hydrocarbon degradation marker genes. To the best of our knowledge, CANT-HYD is the first dedicated tool for annotation of hydrocarbon degradation genes in genomes and metagenomes. We demonstrate the use of CANT-HYD as an exploratory tool by surveying all genomes in GTDB (30,238 bacterial and 1,672 archaeal), as well as several large metagenomic datasets. We uncovered the potential for long-chain alkane degradation in some cyanobacterial genomes and identified widespread potential for aerobic hydrocarbon degradation in global ocean surface waters, supporting a recently discovered marine hydrocarbon cycle.

## DATA AVAILABILITY

The CANT-HYD HMM database and all the custom scripts used in this work can be downloaded through the GitHub (https://github.com/dgittins/CANT-HYD-HydrocarbonBiodegradation) and implemented using any HMM searching tool.

## ACKNOWLEDGEMENTS

We would like to thank Lisa Gieg, Gerrit Voordouw, and Muhe Diao for providing valuable insights into hydrocarbon metabolism. We would also like to thank Dongshan An, Gerrit Voordouw, Daniel Colman, Eric Boyd, Monica Orellana, Avila Viridiana, and Monica Medina for some of the metagenomic data analyzed here. We would like to extend our thanks to Xiaoli Dong for help with access and use of the computational servers. We thank the members of Energy Bioengineering and Geomicrobiology group and countless number of people for their moral support during this “hackathon” that lasted more than a year.

## COFLICT OF INTEREST

The authors declare that the research was conducted in the absence of any commercial, financial, or emotional relationships that could be construed as a potential conflict of interest. The funding agencies are listed in the acknowledgments and did not influence the study design, analyses, outcomes, and publication of this work.

## AUTHOR CONTRIBUTIONS

All authors contributed towards methodology development and editing and reviewing the manuscript. VK, JZ, DG, AC, EB, MAB, AJP, AH, BN, and SB carried out the literature review and sequence data searching. VK, JZ, DG, AC, EB, MAB, AJP, and SB made and validated the HMMs. VK, JZ, DG, AC, EB, and SB wrote the manuscript. VK, JZ, DG, AC, MAB and SB performed genomic and metagenomic data analyses. VK, JZ, AC, and SB made the figures. BN and MS conceived the study. The GitHub archive is created and maintained by DG.

## SUPPLEMENTARY DATA

Supplementary Information – Description of CANT-HYD HMMs

Supplementary Table ST1 – Table of HMMs included in CANT-HYD database Supplementary Table ST2 – Table of experimentally validated sequences used in HMM development

Supplementary Table ST3 – Table of hydrocarbon degrading organisms used for HMM validation

Supplementary Table ST4 – Table of metagenomes tested against CANT-HYD HMM database in case study

Supplementary Figure 1 – Functional homology tree of glycyl radical enzymes Supplementary Figure 2 – Score distribution of AlkB HMM against NCBI nr subset used to determine HMM confidence threshold score

Supplementary Figure 3 – Distribution of CANT-HYD HMM scores among the archaeal representative genomes in GTDB

Supplementary Data SD1 – Tree file for GTDB long-chain monooxygenase analysis

